# iVirus: facilitating new insights in viral ecology with software and community datasets imbedded in a cyberinfrastructure

**DOI:** 10.1101/052597

**Authors:** Benjamin Bolduc, Ken Youens-Clark, Simon Roux, Bonnie L. Hurwitz, Matthew B. Sullivan

**Affiliations:** Department of Microbiology, The Ohio State University, Columbus, OH 43210; Department of Agricultural and Biosystems Engineering, University of Arizona, Tucson, AZ 85721; Department of Civil, Environmental and Geodetic Engineering, The Ohio State University, Columbus, OH 43210

## Abstract

Microbes impact nutrient and energy transformations throughout the world’s ecosystems, yet they do so under viral constraints. In complex communities, viral metagenome (virome) sequencing is transforming our ability to quantify viral diversity and impacts. While some bottlenecks, e.g., few reference genomes and non-quantitative viromics, have been overcome, the void of centralized datasets and specialized tools now prevents viromics from being broadly applied to answer fundamental ecological questions. Here we present iVirus, a community resource that leverages the CyVerse cyberinfrastructure to provide access to viromic tools and datasets. The iVirus Data Commons contains both raw and processed data from 1866 samples and 73 projects derived from global ocean expeditions, as well as existing and legacy public repositories. Through the CyVerse Discovery Environment, users can interrogate these datasets using existing analytical tools (software applications known as “Apps”) for assembly, ORF prediction, and annotation, as well as several new Apps specifically developed for analyzing viromes. Because Apps are web-based and powered by CyVerse super-computing resources, they enable scalable analyses for a broad user base. Finally, a use-case scenario documents how to apply these advances towards new data. This growing iVirus resource should help researchers utilize viromics as yet another tool to elucidate viral roles in nature.

## Viral metagenomics – an ecological tool with increasing impact

Since the first viral metagenomic study was conducted in marine systems over a decade ago (Breitbart *et al.*, 2002), the field has now expanded to include ecological studies of viral communities throughout the oceans globally, as well as diverse lakes and eukaryote-associated samples, including humans (Djikeng *et al.*, 2009; Hannigan *et al.*, 2015; Hurwitz and Sullivan, 2013; Roux *et al.*, 2012; Stern *et al.*, 2012). Highlights of some of the ecological advances enabled by these studies include revealing (i) that virus-encoded ‘auxiliary metabolic genes’ (AMGs) extend far beyond the photosynthesis genes known from cyanobacterial cultures (Hurwitz *et al.*, 2013; Hurwitz, Westveld, *et al.*, 2014; Sharon *et al.*, 2011), (ii) long-term co-evolutionary features between viruses and their microbial hosts in both the human gut (Minot *et al.*, 2013) and the oceans (Hurwitz, Westveld, *et al.*, 2014), and (iii) the ecological drivers of viral community structure throughout the Pacific Ocean (Hurwitz, Westveld, *et al.*, 2014) and global surface oceans (Brum *et al.*, 2015).

Many technological advances have enabled these discoveries – including optimized sampling strategies specific for viruses (reviewed in Duhaime and Sullivan, 2012; Solonenko *et al.*, 2013) and improvements in low-input library preparation methods and decreased sequencing costs (Reuter *et al.*, 2015; Caporaso *et al.*, 2012) – and this has led to a data deluge whereby analytical limitations now represent the major bottleneck for virome-enabled viral ecology as follows. First, the number and size of newly generated metagenomes necessitates large-scale data storage and compute needs that require the development of community-available infrastructures specialized to viral sequence data. Second, the lack of tools for analyzing these large-scale datasets requires development by programmers, who speak a different ‘language’ from researchers generating much of the data. This often results in published code in public repositories that can be difficult to install or use without computational training. Finally, finding and analyzing viral datasets for comparative metagenomics is laborious and time-consuming as raw and processed viral metagenomic datasets are deposited across a diverse array of data repositories, such as Genbank (Benson *et al.*, 1998), EMBL (Kanz *et al.*, 2005), NCBI’s genomes project (Wheeler *et al.*, 2003), Metavir (Roux *et al.*, 2014), and VIROME (Wommack *et al.*, 2012).

Together, these technological limitations impede researchers from applying new tools to their data or leave them dependent on outsourcing data analysis to those unfamiliar with the ecosystems being studied. To enable scalability and accessibility of viral ecology research, we developed iVirus, a collection of software and datasets leveraging the CyVerse cyberinfrastructure (formerly iPlant Collaborative) to provides users with free access to computing, data management and storage, and analysis toolkits (Goff *et al.*, 2011). Briefly, iVirus seeks to collect viral sequence datasets in its Data Commons, adapt pre-existing metagenomic tools as software applications (referred henceforth as *Apps*), and develop new analytical capabilities *within* the CyVerse cyberinfrastructure. Together, these advances help consolidate cutting-edge tools and curated datasets to empower researchers seeking to incorporate viral ecology into their own work.

Here we summarize iVirus’s current capabilities, and invite community feedback, through the protocols.io interface (described later), to allow us to improve iVirus so that it becomes an indispensable tool for ecologists seeking to include viruses in their studies.

## What is CyVerse and how does it help my research?

CyVerse (Goff *et al.*, 2011) is an NSF-funded platform that seeks to bring together biologists and computer scientists to solve ‘big data’ problems in biology. Within the CyVerse cyberinfrastructure, users conduct research by navigating the Discovery Environment (DE) to identify datasets from the Data Commons (next section) and conduct analyses using Apps. Apps are like a computer software program, except that they (i) have been pre-installed, (ii) leverage large-scale CyVerse compute resources, and (iii) can be integrated within a larger data context and workflow. This can improve biological research in multiple ways. First, it helps minimize installation issues and local systems administration needs that often impede biological research. Second, Apps can be linked together to create analytical *workflows* where the output from one App is used as the input for the next, in a linear manner. Because users can select which App they use at each stage in the workflow, the user can copy and update new workflows as new analysis tools emerge. Third, Apps ensure reproducibility and validity of research studies because they can be encoded into versioned Apps, along with raw or processed example datasets directly from the author. CyVerse can also assign digital object identifier (DOI) to Apps or datasets to allow longer term digital preservation and citable referencing in research articles. Finally, because all Apps and their output are tied to the user’s home directory in the CyVerse Discovery Environment, all data are collected in one place. This avoids the common problem of data being scattered among multiple systems (HPC, personal/lab computer, cloud computing) due to system-level requirements for implementing myriad bioinformatics software used for modern metagenomic analyses.

CyVerse Apps can be built by any developer using any source programming language to create community-specific tools. Moreover, the developer can encode hardware or software requirements within the App to lessen the burden on the user in installing and implementing that App. Once an App has been developed it can be shared with the community through the CyVerse cyberinfrastructure by a request to the CyVerse team via a simple public submission form. Once an App is public it can be vetted by the research community via a 5-star rating system and feedback forms. Further, community developers can refine an App by duplicating and modifying it, and then re-publish the new App for recognition and further vetting and use by the community.

For developers, the process of creating an App follows one of two routes. First, Apps can be developed using CyVerse’s API (application program interface) called AGAVE (http://agaveapi.co/) that provides the developer with simple commands to access input data and write logs and results to the CyVerse Data Store. The developer can use the API to specify computational requirements for running code on HPC resources at the Texas Advanced Computer Center (TACC) that are integrated in the CyVerse cyberinfrastructure. This process allows the developer to match the code to HPC compute resources and circumvent difficulties users might experience in installing and re-using the code on different systems. Alternatively, Apps can be deployed using Docker images (www.docker.com), where the code is packaged with additional software dependences and can be run on CyVerse Docker-dedicated servers. Docker’s compute autonomy and portability alone have made Docker images a mainstay for releasing open source code to the user community (Merkel, 2014), in addition to traditional code repositories such as Github (https://github.com). CyVerse extends Docker by allowing developers to deploy Docker images on CyVerse compute resources and attach these images to easy-to-use, web-based Apps in the CyVerse Discovery Environment. This provides developers with a means to publish code in an accessible format to a growing research community and to gain feedback on its utility rather than be drowned in inquiries about installation minutia.

## Centralized viral metagenomic data resources in the iVirus Data Commons

The CyVerse cyberinfrastructure provides a common ecosystem for data, big or small, by providing a mechanism for communities to share data through a Community Data Commons. The iVirus Data Commons leverages these CyVerse resources to make datasets accessible from the Pacific Ocean Virome (POV; Hurwitz and Sullivan, 2013), *Tara* Oceans Virome (TOV; Brum *et al.*, 2015), Southern Ocean Virome (Brum *et al.*, 2016), Virsorter Curated Dataset (Roux, Enault, *et al.*, 2015), and legacy viral datasets from the retired Cyberinfrastructure for Advanced Microbial Ecology Research and Analysis (CAMERA) project (Seshadri *et al.*, 2007). Beyond these, viral data were also mined and hand-curated from Genbank’s Sequence Read Archive (SRA; Benson *et al.*, 1998) and MG-RAST (Wilke *et al.*, 2015). In total, the iVirus Data Commons now contains data from 73 projects, 1 866 samples, and 5.5 billion reads, including contigs assembled from 75 viromes and 121 viral genomes.

## iVirus Apps are Geared to Viral Metagenomics and Community Ecology

iVirus Apps are developed using protocols defined by CyVerse – either through the Agave API or Docker – with focus on those needed for viral metagenomics. One operational goal of iVirus is to collect and deploy the most commonly used tools for viral metagenome pipelines – from raw read processing to assembly and analysis (see overview in Fig. 1, and Use Case Scenario presented in next section). This includes tools for read quality control, assemblers adapted to different input read types and various tools for analyzing assembled viral sequence data.

**Figure 1.**
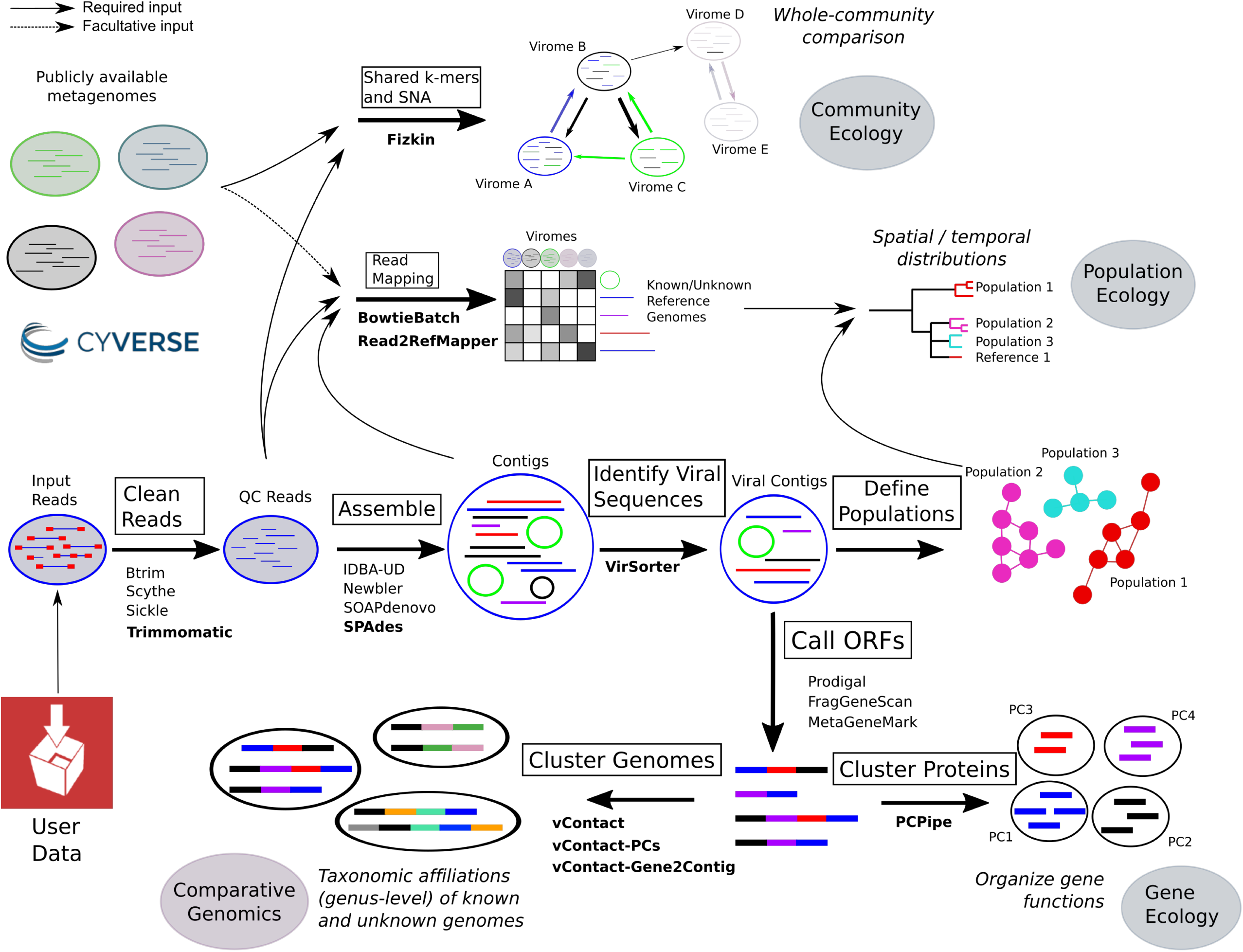
An organizational overview of how a user might leverage iVirus, iMicrobe, and CyVerse Apps to analyze a viral metagenomic dataset. Arrows indicate direction of use case scenario. Processing stages are represented by text in blocks, with bolded text indicating the Apps used in the use case scenario.

Because iVirus exists *within* the CyVerse environment, Apps developed through other community-based CyVerse efforts are also available to iVirus. These include Apps relevant for viral metagenomics such as read pre-filtering for assembly, gene calling, taxonomic identification and sequence alignment tools. This list also includes microbial metagenomic analysis Apps from iMicrobe that can be used to assemble contigs from metagenomes, predict and functionally annotated genes in these contigs, and align them to each other and reference genomes (where available) for comparative genomic analyses. In general, Apps developed for HPC are located in a separate category within CyVerse, under “High Performance Computing” while Docker and non-HPC-enabled Apps are organized into folders appropriate for their “theme” (i.e. iMicrobe, iVirus, Functional Analysis, etc). Because Apps are constantly being updated and developed, a full list of iVirus Apps are maintained at http://ivirus.us/available-tools/, along with computational protocols, App descriptions, relevant articles on tools, and news in protocols.io at http://protocols.io/groups/ivirus

iVirus is a sub-project of iMicrobe and uses open source software developed by that project (http://www.imicrobe.us and http://protocols.io/groups/imicrobe) to query, search, and download data in the iVirus Data Commons from a project specific data website (http://data.ivirus.us/). This web-based resource allows users to perform advanced searches on top of the iVirus Data Commons to discover viral datasets based on related metadata. To better enable search capabilities, iVirus metadata are mapped to the iMicrobe ontology that interconnects existing standards and terminology from the Minimal Information about any (x) Sequence Ontology (MIxS) and community specific ontologies (BCO-DMO, ENVO, CheBI, BCO, OBOE). As such, the location of project specific datasets can be easily discovered and re-used within CyVerse.

Beyond these more generally usable Apps, we have developed several iVirus Apps specifically for viral metagenomic study, with more being added as needs arise and development opportunities become available. In many cases, Apps developed for iVirus and iMicrobe also handle file manipulations, such as compression, separating reads, converting formats, so as to eliminate whenever possible the “minor” details that are often time-consuming and rate-limiting for users.

A brief summary of selected iVirus Apps follows, along with reference to their source tools and how they have already enabled viral ecology where available. A broader analytical pipeline for processing viral metagenomes is overviewed in Figure 1 and described in a User Case Scenario below and at https://www.protocols.io/view/Processing-a-Viral-Metagenome-Using-iVirus-ev3be8n

### PCPipe

This App compares open reading frames (ORFs) from a user-defined dataset to existing viral protein clusters (PCs) as a means to organize proteins derived from viral metagenomics into functional units that can be used as (i) a universal functional diversity metric for viruses, (ii) a scaffold for iterative functional annotations, and (iii) input to ecological comparisons through software such as QIIME, http://qiime.org (Caporaso *et al.*, 2010). This is necessary as viral metagenomes are often dominated by novel sequences, where only 10-20% of reads map to known proteins in reference databases. In contrast, up to 50-70% of reads will typically map to PCs (Hurwitz and Sullivan, 2013). The PCPipe App accepts user-generated ORFs from viral metagenomic assemblies as input, matches them to ORFs in a user-supplied PC database, and then self-clusters the remaining unclustered ORFs to capture the PCs unique to that dataset. Reference sequences from new PCs are annotated using a collection of the non-redundant proteins and associated annotation from the SIMAP database (Rattei *et al.*, 2009). PCs were originally developed for analyzing unknown proteins from the Global Ocean Survey that doubled the known protein universe at the time (GOS; Yooseph *et al.*, 2007). This approach has proved similarly valuable for organizing viral protein sequence space (Brum *et al.*, 2015; Hurwitz and Sullivan, 2013). Such an organizational tool has served as a means to estimate the size of the global virome at a few million proteins (Cesar Ignacio-Espinoza *et al.*, 2013), as well as to make ecological inferences about viral communities with regards to their diversity (Hurwitz and Sullivan, 2013; Brum *et al.*, 2015; Roux *et al.*, 2012), niche differentiating genes (Hurwitz, Brum, *et al.*, 2014) and ecological drivers (Brum *et al.*, 2015).

### VirSorter

This App identifies viral sequences in microbial genomes and metagenomic datasets (Roux, Hallam, *et al.*, 2015). This is necessary as viral genomes are underrepresented in databases – e.g., 92% of 1,659 genome-sequenced phages derive from only 4 of 54 known bacterial phyla (Roux, Enault, *et al.*, 2015). VirSorter can identify diverse viral sequences from microbial datasets, both integrated in the host chromosome and extrachromosomal. Briefly, VirSorter compares a dataset of nucleotide sequences against a user-defined, pre-computed viral database that includes viral sequences from RefSeq and (if desired) contigs assembled from viral metagenomes. The comparison also takes into account viral hallmark genes, as well as statistical enrichment of viral genes, depletion in hits to the PFAM database, and strand bias. VirSorter output includes a summary file with “confidence” categories for each identified sequence, as well as predicted proteins, PFAM domain hits, suspected circular sequences and metrics files. This tool is powerful and highly scalable – its first application was to nearly 15 000 publically available archaeal and bacterial genomes, where VirSorter identified 12 498 new host-associated viruses and their genomes, which augmented publicly available viral genome reference datasets approximately 10-fold (Roux, Hallam, *et al.*, 2015). Further, VirSorter scales to handle contigs derived from metagenomic datasets (Roux, Enault, *et al.*, 2015). VirSorter has since been used to identify viruses out of boiling hot springs in Yellowstone National Park (Munson-McGee *et al.*, 2015), the *Tara* Oceans Viromes (Lima-Mendez *et al.*, 2015), and hypersaline environments in the Atacama Desert (Crits-Christoph *et al.*, 2016).

### vContact

This App assigns contigs to taxonomic groups using the presence or absence of shared PCs along the length of the contig. This is critical as viruses lack a universal gene marker (Edwards and Rohwer, 2005) and less than 0.1% of viruses in natural environments are represented in public databases (Brum *et al.*, 2015), which necessitates new approaches to taxonomically classify surveyed viral genomes. Inspired by algorithms to detect prophage in microbial genomes (Lima-Mendez *et al.*, 2008), vContact clusters contigs by their PC profiles (note: see preferred method for PC generation as vContact-PCs, but the user could generate PCs however they prefer). Reference sequences and their taxonomic lineages can be seeded within the analysis to improve clustering and taxonomic predictions. The vContact-generated network can be mined for its contig clusters (VCs: viral clusters), which roughly correspond to subfamily level viral taxonomy. This approach has been used to organize nearly all (99.3%) of the 12,498 new viral sequences identified from publicly available microbial genomes into 614 ‘viral clusters’, which represent approximately genus-level groupings (Roux, Hallam, *et al.*, 2015). vContact can also incorporate annotations associated with contig and PCs, allowing users to examine the relationship of any annotated contig/PC in context of its vContact cluster.

### vContact-PCs

This App serves as a companion tool for vContact to generate PCs using a Markov clustering algorithm (MCL; Enright *et al.*, 2002). Users provide a BLASTP file of an all-against-all protein comparison, and vContact-PCs parses the BLAST file and applies MCL against its similarity scores. vContact-PCs then exports files formatted for use with vContact.

### Fizkin

This App performs Bayesian network analyses based on the amount of shared sequence content in viromes and relevant environmental metadata. Specifically, the App randomly subsets 300K reads (or the lowest common denominator for reads in viromes) from up to 15 viromes and performs a pairwise all-vs-all kmer-based sequence comparison between all virome pairs as previously described (Hurwitz, Westveld, *et al.*, 2014). The kmer search in Fizkin is implemented in Jellyfish (Marcais and Kingsford, 2011) and is used to rapidly generate a matrix of shared sequence counts between each virome pair. This matrix is then used as input into a Bayesian network analysis (Hoff, 2005). The user can also input environmental metadata (continuous or discrete measurements, or latitude and longitude) that are used as part of the analysis to determine which environmental factors are significant in defining the structure of the network. Output includes: (i) a table indicating which environmental factors are significant and (ii) a social network graph to visually represent the distance between viromes where statistical samples are taken from the marginal posterior distributions, and samples names are placed at the posterior mean. This type of social network analysis has been used to evaluate viral community structure and ecological drivers (Hurwitz, Westveld, *et al.*, 2014), as well as to quantify lysogeny through comparative analysis of experimentally induced and non-induced viromes (Brum *et al.*, 2016).

### BatchBowtie

This App runs bowtie2 on any number of reads files within a directory the user selects. Read recruitment against reference sequences (i.e viral genomes) can be employed as a means to visualize spatial or temporal distributions of genomes via the reads relative abundance across samples (Brum *et al.*, 2015). The App additionally offers the ability to convert between interleaved and non-interleaved fastq files, compressed files, and can generate SAM and BAM-formatted outputs (typically used by Read2RefMapper).

### Read2RefMapper

This App consumes BAM alignment files (i.e. from BatchBowtie), generating coverage tables and relative abundance plots – useful for identifying the abundance of reads against a set of reference sequences. Users can select a variety of filtering options based on the percent of read *and* reference covered (i.e. 75% of a reference sequence must be covered to be considered “present”), alignment identities, as well as numerous coverage calculations. If users provide a file with the size of each metagenome, Read2RefMapper will also normalize the coverage between samples.

## Finding and using iVirus: A use case scenario ‘live’ at protocols.io

Taken together – the Apps mentioned above, in addition to the Apps already available in CyVerse – can be used to process a viral metagenome from “raw” sequence to minimally characterized viral assemblies. The guide(s) for this *and other* examples are available at protocols.io: https://www.protocols.io/view/Processing-a-Viral-Metagenome-Using-iVirus-ev3be8n. These protocols are organized as collections, and available within the iVirus and iMicrobe groups at protocols.io, at https://www.protocols.io/groups/ivirus and https://www.protocols.io/groups/imicrobe. These groups serve as a centralized location that offers additional documentation, feedback, as well as citations using these tools and protocols. We have utilized protocols.io here so as to keep evolving processing steps up-to-date, as well as include images and annotations for each step. Further, protocols.io is ideal for obtaining community feedback as it provides users the opportunity to ask questions and/or interact with the protocol’s author through a simple to user interface.

The use case scenario starts with test data, which are reads from publicly available Ocean Sampling Day 2014 samples (https://github.com/MicroB3-IS/osd-analysis/wiki/Guide-to-OSD-2014-data), a subset of which are already available on CyVerse’s data store. Next a user must first register for a *free* account in CyVerse (http://user.iplantcollaborative.org/) and then proceed through the process using available iVirus Apps summarized in Fig. 1. While only a few Apps are highlighted for each step, there are wide selections of choices for Apps available to the user. Example data is provided for each step in the iVirus Data Commons found under the /iplant/home/shared/iVirus/ExampleData/ (https://de.iplantcollaborative.org/de/?type=data&folder=/iplant/home/shared/iVirus/ExampleData) folder within the CyVerse Discovery Environment. Output from each stage in this scenario is used as input for the next, though users can test any step individually, as each stage is organized separately and contains its own folders with inputs and outputs to help users identify which files are associated with each step.

### <u>Step 1</u>: Upload Read Data

Before processing can begin, reads to be analyzed must be uploaded to CyVerse’s Data Store in the user’s account. Small files can be uploaded from the user’s computer through the Discovery Environment’s (DE) upload menu, with larger files transferrable through popular SFTP software, such as Cyberduck (https://cyberduck.io) and iRODS (http://irods.org). Data uploaded to a user’s CyVerse home directory are private and accessible only to the user until they grant access to collaborators (via private invitation) or publish their data to the larger CyVerse community by transferring files to a shared Community folder. Users can also analyze or leverage growing publicly available datasets at iVirus in the Data Commons as previously described. Use case read files are already uploaded to the iVirus Data Commons.

### <u>Step 2</u>: Quality Control (QC) of Read Data using Trimmomatic

Protocol available at protocols.io: https://www.protocols.io/view/Quality-Control-of-Reads-Using-Trimmomatic-Cyverse-ewbbfan

Once raw read data has been uploaded, reads need to be quality filtered to ensure high quality reads for assembly. FastQC (http://www.bioinformatics.babraham.ac.uk/projects/fastqc/) is one such App already available through CyVerse, and provides a visualization of read lengths, quality scores, duplicate reads, N and GC content of raw reads that can be used to determine the appropriate parameters for quality control. Once quality-filtering parameters have been determined for a given sequencing run via FastQC, reads files can be trimmed and quality controlled by using the Trimmomatic App.

### <u>Step 3</u>: Assembly of QC Read Data using SPAdes

Protocol available at protocols.io: https://www.protocols.io/view/Assembling-Viral-Metagenomic-Data-with-SPAdes-Cyve-evzbe76

Following QC, reads are then assembled using one of the assemblers available in CyVerse. Most frequently, assembler selection is based on read type (Sanger, 454, Illumina, PacBio, etc) and to a lesser extent, its performance for a particular sample type. IDBA-UD, SOAPDenovo, Trinity and SPAdes are available for viral metagenomic assembly. Some assemblers have high memory variants for larger data sets, and should only be used when the standard versions fail to assemble.

### <u>Step 4</u>: Identification of Viral Sequences from Assembled Data using VirSorter

Protocol available at protocols.io: https://www.protocols.io/view/Identifying-Viral-Sequences-Using-VirSorter-Cyvers-ev2be8e

To identify viral sequences from the contigs file, VirSorter is used. Generally, contigs larger than 3-kb can be successfully used as input – and can be single cell genomes, microbial or viral metagenomes, fragmented or complete genomes.

### <u>Step 5</u>: Characterizing Viral Sequence Data through Protein Clustering with PCPipe and vContact

Protocol available at protocols.io: https://www.protocols.io/view/Preparing-Data-for-vContact-from-Proteins-Cyverse-ev7be9n

Protocol available at protocols.io: https://www.protocols.io/view/Applying-vContact-to-Viral-Sequences-and-Visualizi-ev8be9w Large-scale characterization of viral genomic data remains one of the most daunting challenges in viral ecology. A relatively recent method of analyzing complex viral data is through *organizing* viral sequence space using PCs (protein clusters), reducing the problems associated with data complexity as a byproduct. Regardless of the type of analysis, iVirus has access to a number of tools to characterize viral data.

## Concluding Remarks

While viruses are increasingly recognized for their roles in microbial-dominated ecosystems, they remain understudied, particularly due to challenges stemming from the lack of centralized viral metagenomic resources. iVirus offers a community-focused resource, built on the CyVerse cyberinfrastructure and designed to directly address the challenges of viral ecology in the era of next-generation sequencing, high performance computing and big data analytics. This is done through 1) leveraging CyVerse’s Data Store to provide large data storage capacity and a centralized location for collecting data, 2) developing Apps, or software applications designed to take advantage of HPC resources that require limited bioinformatics training on part of the researcher, 3) collecting viral datasets in the iVirus Data Commons to provide a centralized location for discovering datasets via environmental metadata and collaborating within the field, and 4) positioning these resources to maximize community exposure and feedback through extensive and ‘live’ documentation at protocols.io.

## Acknowledgements

The authors wish to thank Joanne B. Emerson, Dean R. Vik, Gareth G. Trubl, Consuelo Gazitua, Pilar Manrique, and Jacob H. Munson-McGee for their helpful feedback in designing community-minded tools; Matthew Vaughn from the Texas Advanced Computer Center (TACC) and Nirav Merchant from the BIO5 Institute whose assistance in App development, troubleshooting and making Apps publicly available were invaluable. We thank Lenny Teytelman, Lori Kindler, and Alexei Stoliartchouk for their agile development of protocols.io to accommodate requests for the iVirus group. The development of iVirus and this publication was partially funded in part by Gordon and Betty Moore Foundation grants GBMF4491, GBMF4733, GBMF3305, GBMF3790, by the US Department of Energy Office of Biological and Environmental Research under the Genomic Science program (Award DE-SC0010580) and by a TRIF award to the University of Arizona Ecosystem Genomics Initiative.

